# A miniaturized ultrasound transducer for monitoring full-mouth oral health: a preliminary study

**DOI:** 10.1101/2022.03.31.486598

**Authors:** Baiyan Qi, Ali Hariri, Reza K. Nezhad, Lei Fu, Yi Li, Zhicheng Jin, Wonjun Yim, Tengyu He, Yong Cheng, Jiajing Zhou, Jesse V. Jokerst

**Affiliations:** Materials Science and Engineering Program, University of California San Diego, La Jolla, California 92093, USA; StyloSonic LLC, Lake Forest, California 92630, USA; Department of Nanoengineering, University of California San Diego, La Jolla, California 92093, USA; Department of Radiology, University of California San Diego, La Jolla, California 92093, USA

**Keywords:** periodontitis, gingiva, ultrasonography, ultrasonic imaging

## Abstract

**Objective:** To customize a miniaturized ultrasound transducer to access full-mouth B-mode, color Doppler and spectral Doppler imaging for monitoring oral health.

**Methods:** A customized periodontal ultrasound transducer SS-19-128 (19 MHz, 128 channels) with 1.8 cm wide and 1 cm thick was developed and connected to a data acquisition (DAQ) system. B-mode, color Doppler, and spectral Doppler data could all be collected with SS-19-128. The imaging resolution and penetration capacity of SS-19-128 were characterized on phantoms. Five human subjects were recruited to demonstrate B-mode and Doppler imaging by SS-19-128. Gingival thickness was measured on 11 swine teeth by SS-19-128 for comparison to conventional transgingival probing via Bland-Altman analysis and Pearson correlation.

**Results:** The axial and lateral spatial resolution at 5.5 mm depth is 102.1 μm and 142.9 μm, respectively. The penetration depth in a tissue-mimicking phantom is over 30 mm. *In vivo* B-mode imaging of all 28 teeth was demonstrated on one human subject, and imaging of tooth #18 was accessed on five human subjects. Gingival thickness measurement compared with transgingival probing showed a bias of −0.015 mm and SD of 0.031 mm, and a r = 0.9235 (P<0.0001) correlation. *In vivo* color and spectral Doppler imaging of the supraperiosteal artery in human gingiva was performed to generate hemodynamic information.

**Conclusions:** The small size of SS-19-128 offers important advantages over existing technology—more specifically, whole-mouth scanning/charting reminiscent of radiography. This is nearly a two-fold increase in the number of teeth that can be assessed versus existing transducers.

## INTRODUCTION

Periodontitis can lead to tooth decay, tooth loss,(1) and even systemic cardiovascular disease;(2) 50% of Americans have periodontitis.(3) Thus, monitoring anatomical and hemodynamic oral health is important. Radiography and periodontal probing are current standard methods to diagnose periodontal anatomy, and laser Doppler flowmetry has been used for hemodynamic oral research.(4) However, existing tools suffer from major limitations in diagnosing and monitoring periodontitis.

Radiography provides excellent sensitivity to hard tissues, but uses ionizing radiation and is limited in stratifying disease in soft tissues. Periodontal probing is time-consuming and painful for patients, and inter-operator variation in probing can be larger than 40%:(5) This leads to large measurement errors. Laser Doppler flowmetry can only detect movement of erythrocytes in a small tissue volume (about 1 mm^3^), which prevents it from analyzing blood flow in a larger region of interest.(6) Furthermore, laser Doppler flowmetry suffers from poor reproducibility because a minimal displacement of the probe can cause the change of the investigated area due to the dense gingival vascular network.(7) Compared with laser Doppler flowmetry, Doppler ultrasound measurement offers real-time diagnosis as well as high reproducibility that allows monitoring oral microcirculation changes over time; metal constructions and/or implants do not limit studies of blood flow in the gingiva or tooth pulp.(8)

Previously, we and others (9–14) have reported that ultrasound imaging has major value in oral health and can improve the limitations of conventional methods.(15, 16) Ultrasound offers visualization of anatomical features including the alveolar bone (AB), alveolar bone crest (ABC), cementoenamel junction (CEJ), gingival margin (GM), gingival thickness, etc. in humans, *ex vivo* swine jaws, and cadavers.(11, 17–20) Machine and deep learning can improve image analysis.(12, 21, 22) Other studies also improve transducer design, hardware, and coupling materials for ultrasound imaging.(23, 24) However, clinical translation and validation of ultrasound in oral health remains difficult because existing transducers are too bulky to access posterior teeth. While many compact and endoscopic ultrasound transducers are available commercially, none offer the high frequency needed to resolve small dental feature sizes.

Here, we characterized a custom-built 128-element, 19-MHz transducer (SS-19-128) for periodontal imaging including all posterior teeth. The 1.8 cm × 1 cm form factor ensures access to the entire oral cavity including all pre-molars and molars. First, we measured the transducer’s axial/lateral resolution and imaging depth. Then, both human and cadaver swine were imaged to identify landmarks in the oral anatomy. We then measured the gingival thickness in a swine model and compared image-based metrics to gold standard techniques (i.e., transgingival probing). We then conducted a whole-mouth scan and evaluated real-time blood flow in human gingiva.

## MATERIALS AND METHODS

### Equipment

This work used a customized transducer (SS-19-128, StyloSonic, Lake Forest, USA). SS-19-128 has a 128-element linear array with a center frequency of 19 MHz and an average −6 dB bandwidth of 48.9%. The length of the ultrasound active area is 10.24 mm with an element pitch size of 78 μm. The transducer is housed inside a toothbrush-shaped handpiece (1.8 cm × 1 cm) to access the entire oral cavity. SS-19-128 has a side-facing view within the housing to ensure proper positioning with the gingiva and tooth surface. Disposable Tegaderm films (3M, Minnesota, USA) were applied as transducer sleeves for sterility when doing human scans.

SS-19-128 was connected to a data acquisition (DAQ) system (Vantage 256, Verasonics, Inc., Kirkland, USA) via a UTA 260D adapter. The DAQ provided a 5.6-V power supply and controlled electrical excitation time delay for each channel. The DAQ sampled the radiofrequency data from all 128 channels and processed ultrasound images. This system supports a frequency range of 2 to 42 MHz; 14-bit A/D converters with a programmable sample rate up to 62.5 MHz and can image up to 100,000 frames/second.

### Human Subjects Recruitment

The study protocol with human subjects was approved by the UCSD Institutional Review Boards (project 170912) and was conducted according to the ethical standards set forth by the IRB and the Helsinki Declaration of 1975. Five healthy adults with good oral hygiene were enrolled. All subjects gave informed consent.

### Preparation of phantoms

To characterize imaging resolution, two 30-µm nichrome wires were placed in water and held 5.5 mm and 13.5 mm from the surface of SS-19-128. A tissue mimicking phantom we previously reported was used to demonstrate the penetration depth of SS-19-128.(25)

### B-Mode Ultrasound Imaging

Three different modes of beamforming were used to process B-mode ultrasound images: (i) plane wave, (ii) coherent compounding, and (iii) synthetic aperture.(26) In plane wave imaging, all channels of the transducer were excited spontaneously to generate one plane wave with no beam tilting; a B-mode image was reconstructed from the ultrasound pulse-echo signal.(27) In coherent compounding imaging, a time delay between channels was applied by the DAQ system.(28) Seven plane waves with different tilting angles were then generated in a row, and one image frame was reconstructed by averaging the images obtained with all tilted plane waves. In synthetic aperture imaging, the transducer channels were divided into many subgroups. In each ultrasound image frame, the subgroups were sequentially excited and swept from one side to the end along the transducer array.(27) The reconstruction process compensated for the phase differences and then added up the pulse-echo signals from all channels.

### Color and Spectral Doppler Imaging

Color Doppler imaging is a combination of B-mode gray-scale imaging and blood flow velocity measurement by mapping Doppler frequency shift of pulsed ultrasound waves. Based on the Doppler effect, ultrasound waves reflected by blood that moves towards the ultrasound transducer are increased in frequency, and vice versa.(29) In the region of interest, increased frequency is displayed in red, and decreased frequency is in blue.

The received Doppler signal contains Doppler shifts at different frequencies. These shift frequencies were decoded by a fast Fourier transform and displayed as a spectrum: The vertical axis is the Doppler frequency shift, and the horizontal axis is time. The amplitude of the Doppler shift frequency is presented as brightness. The power of the Doppler frequency shift signal is proportional to the number of blood cells per unit volume. A bright spot on the Doppler spectrum means that a strong Doppler frequency shift amplitude is received at that instant of time, which suggests that there are more scattering blood cells moving at that speed.

### Gingival Thickness Measurement

Swine jaws were purchased from the local butcher and sliced with a bandsaw to size. They were immobilized in water for ultrasound coupling for imaging. Values of gingival thickness were measured by SS-19-128 ultrasound imaging and a 28-gauge needle with calipers (transgingival probing; gold standard method).(30–32) The gingival thickness measurement point was defined as 2 mm below the GM. In the ultrasound imaging method, the gingival thickness was obtained by measuring the distance between the gingival surface and the tooth surface via ImageJ software.(33) In the clinical periodontal probing method, the needle was inserted into the gingiva perpendicularly until contact with the tooth.(32) A line was marked on the needle, and the distance between the marked line and the tip of the needle was measured with a 0.1 mm precision caliper.

### Statistical analysis

Bland-Altman plot and Pearson correlation were analyzed by GraphPad Prism 9 (GraphPad Software, San Diego, California USA, www.graphpad.com) to compare the gingival thickness measurement on 11 swine teeth via SS-19-128 and periodontal probing method.

## RESULTS

The periodontal ultrasonography system (**Fig. 1a, b**) includes SS-19-128 and a DAQ system that provides power supply, electrical excitement control, radiofrequency signal decoding, and real-time ultrasound imaging display. SS-19-128 was connected with a universal transducer adapter suitable for interfacing with the DAQ (**Fig. 1c**). The side-facing SS-19-128 was designed as a toothbrush shape to image posterior teeth (**Fig. 1d**). The accessible teeth imaging range between the recent work (13) and SS-19-128 was compared (**Fig. 1e**).

**Figure 1.**
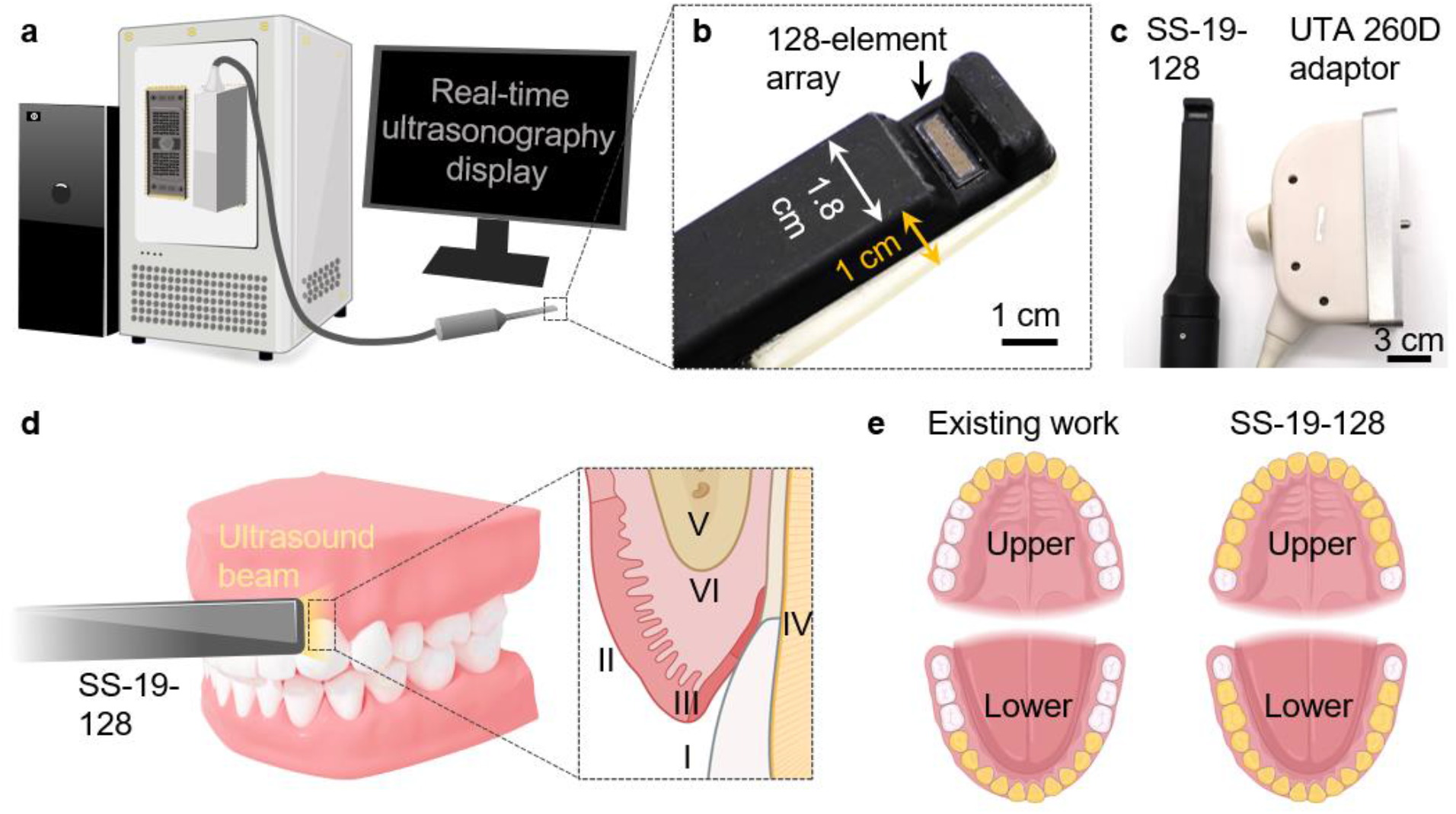
Overview of the customized transducer and periodontal imaging. (**a**) SS-19-128 was connected to the DAQ system for real-time ultrasonography display. (**b**) Photograph of the 128-channel array located in the 3-D printed housing. (**c**) Lower magnification photograph of the entire handpiece and the UTA 260D adaptor to the DAQ. (**d**) Schematics of the side-facing SS-19-128 being applied to human tooth. Six anatomical features were determined: I – tooth surface; II – gingiva; III – GM; IV – CEJ; V – AB; VI – ABC. (**e**) Comparison of accessible imaging range between our prior work (13) and SS-19-128 with size miniaturization. Accessible teeth were marked as yellow.

### Evaluation of Performance Using Phantoms

The axial and lateral resolutions of SS-19-128 were characterized using line spread function (34) to nichrome wire images (**Fig. 2b, c**). The axial resolutions of plane wave and coherent compounding modes were the same (102.1 μm), which was 9.2% higher than the synthetic aperture mode (112.5 μm).

**Figure 2.**
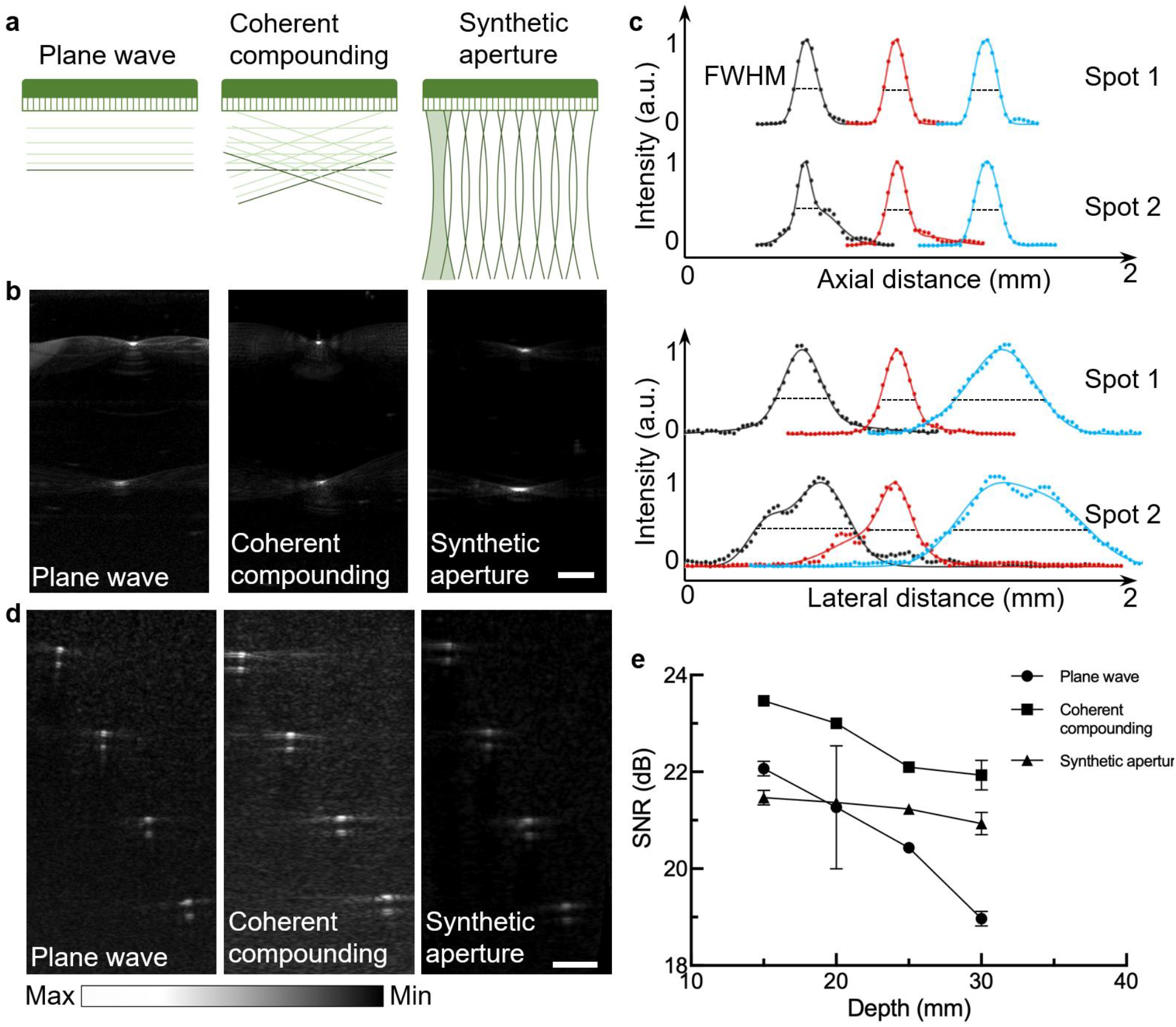
Resolution and penetration characterization. (**a**) Illustration of the three beamforming modes used here for ultrasound B-mode imaging. (**b**) Ultrasound imaging of nichrome wires at 5.5 and 13.5 mm with three beamforming modes. (**c**) Axial and lateral resolution characterization at 5.5 and 13.5 mm. (**d**) Ultrasound imaging of tubes buried in tissue-mimicking phantom at 15, 20, 25, 30 mm with three beamforming modes. (**e**) SNR vs depth curve under three beamforming modes.

The lateral resolution of the wire at 5.5 mm from the transducer was 142.9 μm in coherent compounding mode—this was 36.4% higher than plane wave mode (224.6 μm) and 58.8% higher than synthetic aperture mode (347.1 μm).

We tested the penetration depth of SS-19-128 with a tissue-mimicking phantom in which four polytetrafluoroethylene tubes were embedded at 15 mm, 20 mm, 25 mm and 30 mm (**Fig. 2d**). The signal-to-noise ratio (SNR) of the 30-mm-deep tube was 19 dB, 22 dB, and 21 dB under plane wave, coherent compounding, and synthetic aperture modes, respectively. The SNR *versus* depth under three beamforming modes is further quantitated in **Fig. 2e**.

### Periodontal Imaging of Five Human Subjects

This human subject work involved five volunteers. On one subject, 28 teeth were accessed and imaged by SS-19-128 (#2 to #15 and #18 to #31; four wisdom teeth had been extracted; **Fig. 3a**). A high magnification image of one representative tooth #13 is shown in **Fig. 3b**, and six anatomical features were marked. **Fig. 3c** shows the corresponding panoramic X-ray image of the same human subject. To demonstrate the reproducibility the customized transducer to access all teeth, tooth #18—the most posterior mandibular left molar—was imaged in five healthy subjects with clearly visualized anatomical features (**Fig. 3d**).

**Figure 3.**
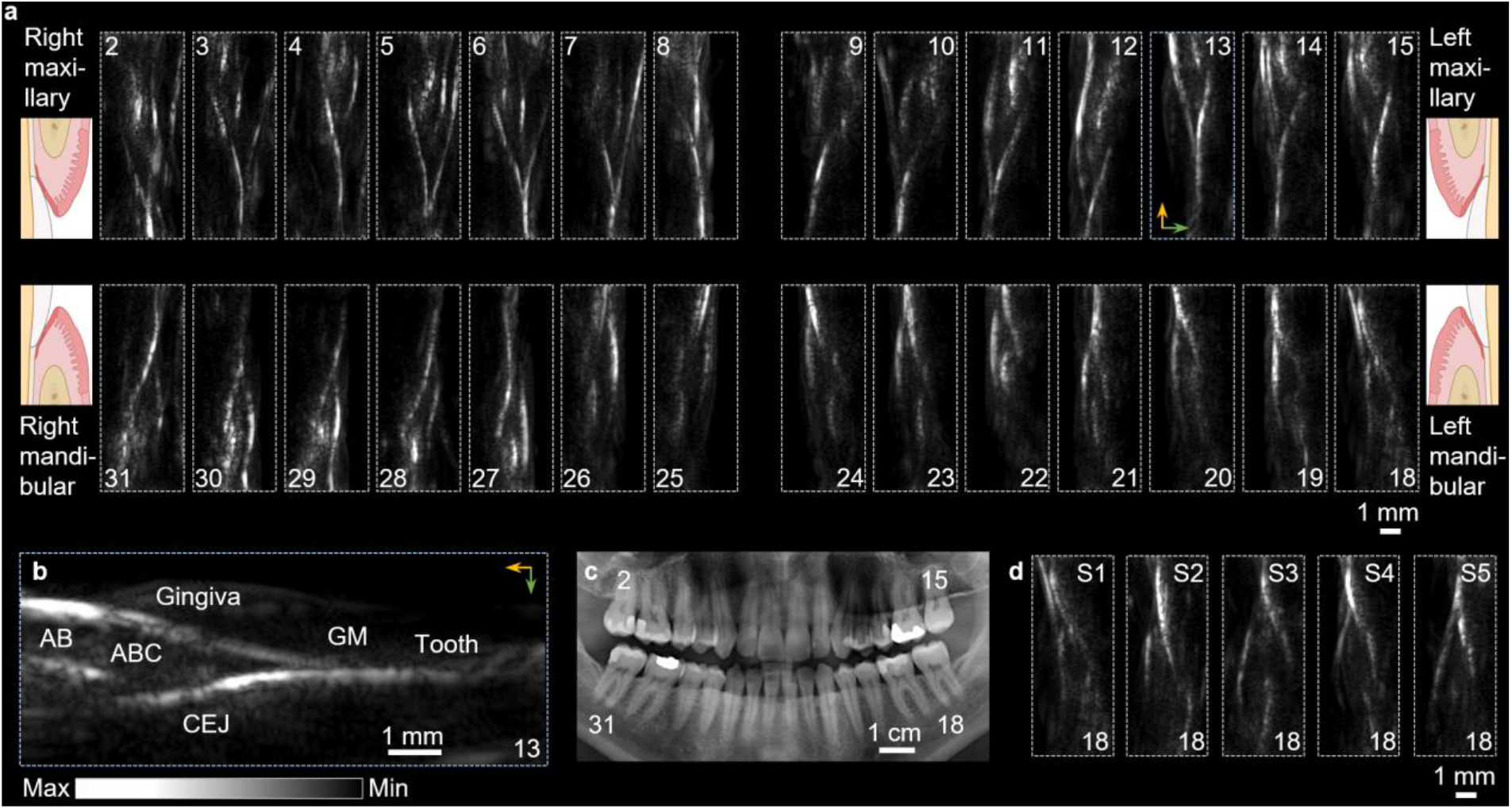
*In vivo* imaging on human teeth. (**a**) Full-mouth scan of 28 teeth in a healthy human subject. Corresponding schematics showed the orientation of imaging of four quadrants, respectively. (**b**) A representative tooth #13 from the full-mouth imaging. Anatomical structures including tooth, gingiva, GM, CEJ, AB, and ABC are marked in the tooth imaging. Image orientation was marked by arrows. (**c**) Full-mouth panoramic X-ray image of the same human subject. (**d**) Image of tooth #18 (most posterior molar other than wisdom tooth) of five different healthy subjects (labelled as S1-S5). Image orientation is consistent with the left mandibular quadrant in (a).

### Evaluation of Gingival Thickness Measurement Using 11 Swine Teeth

The gingival thickness was measured on 11 swine teeth with SS-19-128 (**Fig. 4a**) as well as with a periodontal probing needle. The gingival thickness measured with SS-19-128 was termed as SS-GT; while the one measured with transgingival probing was termed as TP-GT. The mean SS-GT was 1.34 ± 0.08 mm and the mean TP-GT was 1.32 ± 0.07 mm. Gingival thickness measurements correlated well (**Fig. 4b**) between the two methods (r = 0.9235, p < 0.0001). The Bland-Altman analysis (**Fig. 4c**) for gingival thickness revealed a bias of −0.015 mm and SD of 0.031 mm, and a 95% limits of agreement from −0.077 mm to 0.046 mm.

**Figure 4.**
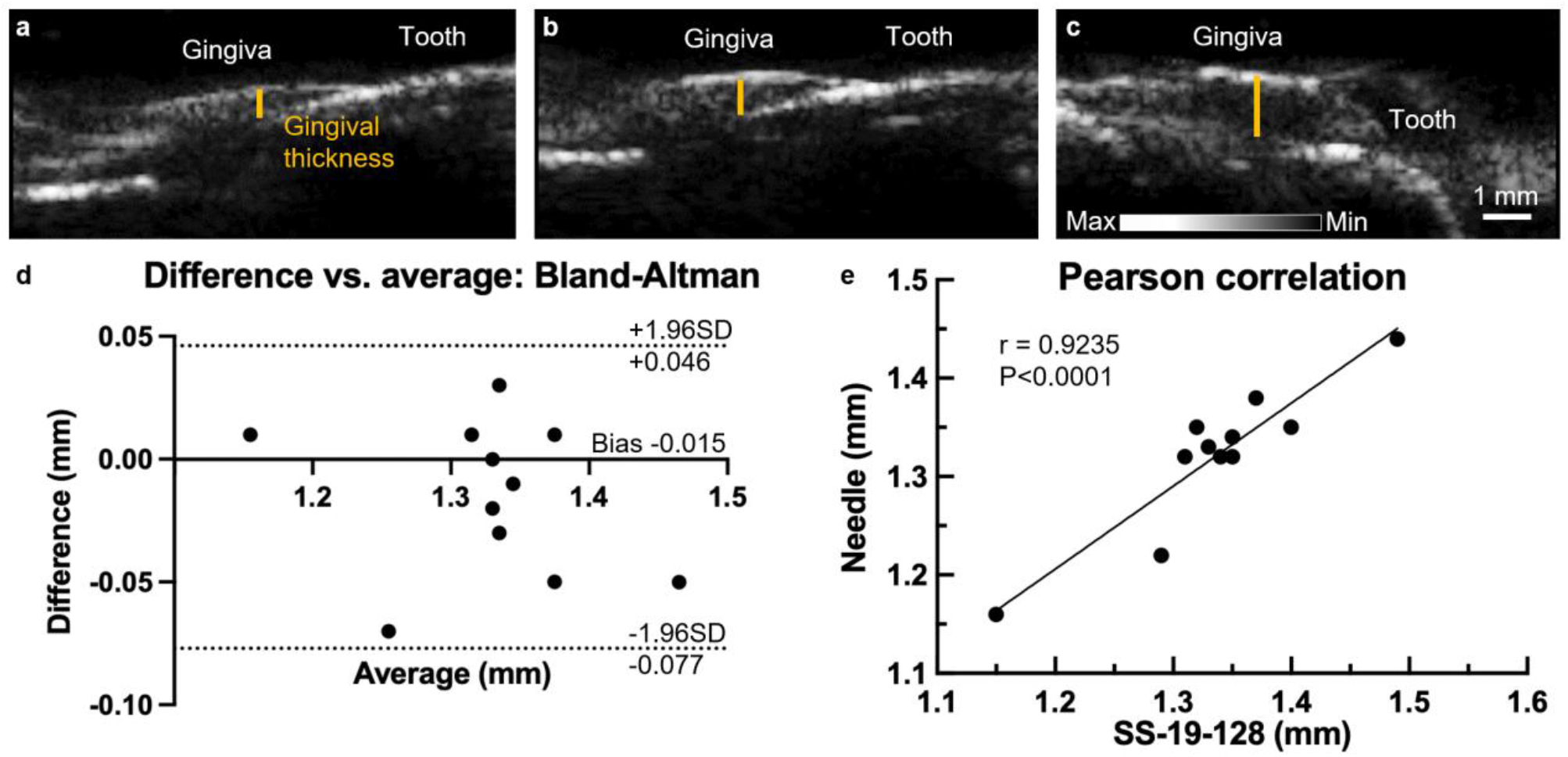
Gingival thickness measurement on swine jaws and statistical analysis. SS-19-128 transducer imaged swine teeth with gingival thickness including small (**a**), medium (**b**), and larger gingival thickness (**c**); yellow line indicates approximate area of measurement. (**d**) Bland-Altman plot comparing ultrasound imaging and needle measurements indicating a −0.15 mm bias between two measurements. (**e**) Pearson correlation between ultrasound imaging and needle measurement. Correlation coefficient r = 0.9235 (P < 0.0001) showing a high correlation relationship between two measurements.

### Evaluation of Gingival Blood flow Measurement of Human

Here, color Doppler and spectral Doppler imaging were established in SS-19-128 to monitor hemodynamic signals of blood supply of supraperiosteal artery in gingiva of a human subject.

Color and spectral Doppler imaging was processed on human radial artery on wrist and cadaver swine jaw via SS-19-128 to serve as positive (blood flow existed) and negative (no blood flow) control, respectively. The red color inside the radial artery indicated the Doppler frequency shift of the received radiofrequency signal is <0, which means the blood flow direction was towards the transducer (**Fig. 5a**). By decoding the radiofrequency signals of the center of radial artery by fast Fourier transform, the spectral Doppler signals along time were processed and displayed in real-time (**Fig. 5b**). In the cadaver swine jaw, no Doppler signals were detected because there was no blood flow (**Fig. 5c, d**). Color and spectral Doppler were then performed on human tooth #28 by SS-19-128. Blood flow of supraperiosteal artery in gingiva was imaged using color Doppler, and the red color showed that blood was moving towards the transducer (**Fig. 5e**). The decoded spectrum of the artery was then performed (**Fig. 5f**).

**Figure 5.**
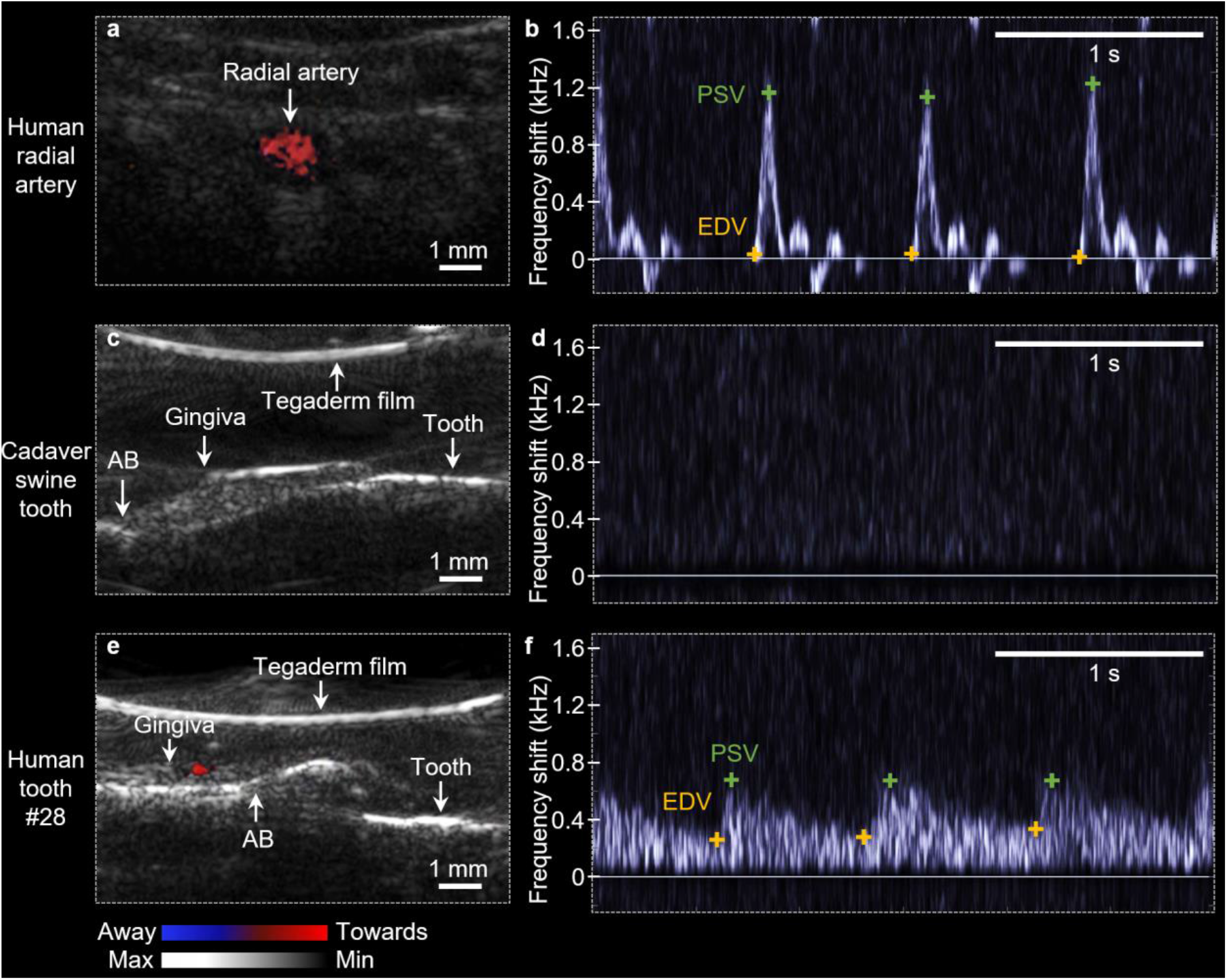
Doppler imaging on human tooth. (**a**) Color Doppler image on human radial artery (positive control). Red color indicates the blood moved towards the transducer. (**b**) Spectral Doppler of the radial artery. PSV: peak systolic velocity; EDV: end diastolic velocity. (**c**) Color and (**d**) spectral Doppler images on cadaver swine tooth (negative control). (**e**) Color and (**f**) spectral Doppler images on human tooth #28.

## DISCUSSION

The cheeks restrict the movement and positioning of larger transducers or transducers with existing front-facing designs, and thus their accessible range is limited to anterior teeth. To date, The Michigan group reported ultrasound imaging in maxillary incisors and the mandibular second pre-molar by transducer L25–8;(35) the total number of accessible teeth could be estimated as 14 with that transducer. Fu et al. successfully accessed human teeth #5 and #29 by transducer CL15-7 for 18 total accessible teeth.(13) In contrast, SS-19-128 can perform a full-mouth scan, and its accessible range was increased to all 28 teeth (wisdom teeth were extracted), which is 55.6% higher compared with the existing work.

In ultrasound imaging, higher frequency provides high resolution but suffers from lower penetration depth. Fortunately, periodontal ultrasound rarely requires deep imaging (i.e., maximum is about 1 cm from tooth surface).(36) However, the size of the oral features (CEJ, gingival thickness, etc.) are often on the sub-millimeter scale, which requires high-resolution imaging.(37) Thus, there is value in developing transducers with higher frequencies because they offer better resolution. It is known that axial resolution is equal to spatial pulse length (SPL) divided by 2,(38) and SPL is proportional to ultrasound wavelength and pulse cycle number.(39) The wavelength and pulse cycle number were the same in three beamforming modes, and thus the values should be identical; we noticed a ∼10% difference perhaps due to subtle differences in the positioning of the phantom relative to the transducer. In the lateral direction, the artifacts in coherent compounding mode were compressed by averaging radiofrequency signals from plane waves with seven tilting angles.(40) Our goal here was high resolution for the small oral features, and thus coherent compounding beamforming satisfied the resolution requirement best.

To compare with existing periodontal transducers, the axial resolution of L25-8 (24 MHz) (35) was 64 μm, 37% higher than SS-19-128 (102.1 μm) because of its higher center frequency. Although the axial resolution of SS-19-128 was relatively lower, it was sufficient for periodontal imaging because features sizes are on the millimeter scale (e.g., median gingival thickness =1.1 mm).(41) The axial and lateral resolution of the Philips CL15-7 (9 MHz) (13) was 210 μm and 196 μm, respectively. The axial resolution of CL15-7 was 106% lower than SS-19-128 because of its lower center frequency. The lateral resolution was not fair for comparison because the distance between imaging targets and the transducer CL15-7 was not mentioned in this work. Additional comparison of parameters that define the performance of an ultrasound transducer is included in **Table 1**.

**Table 1.**
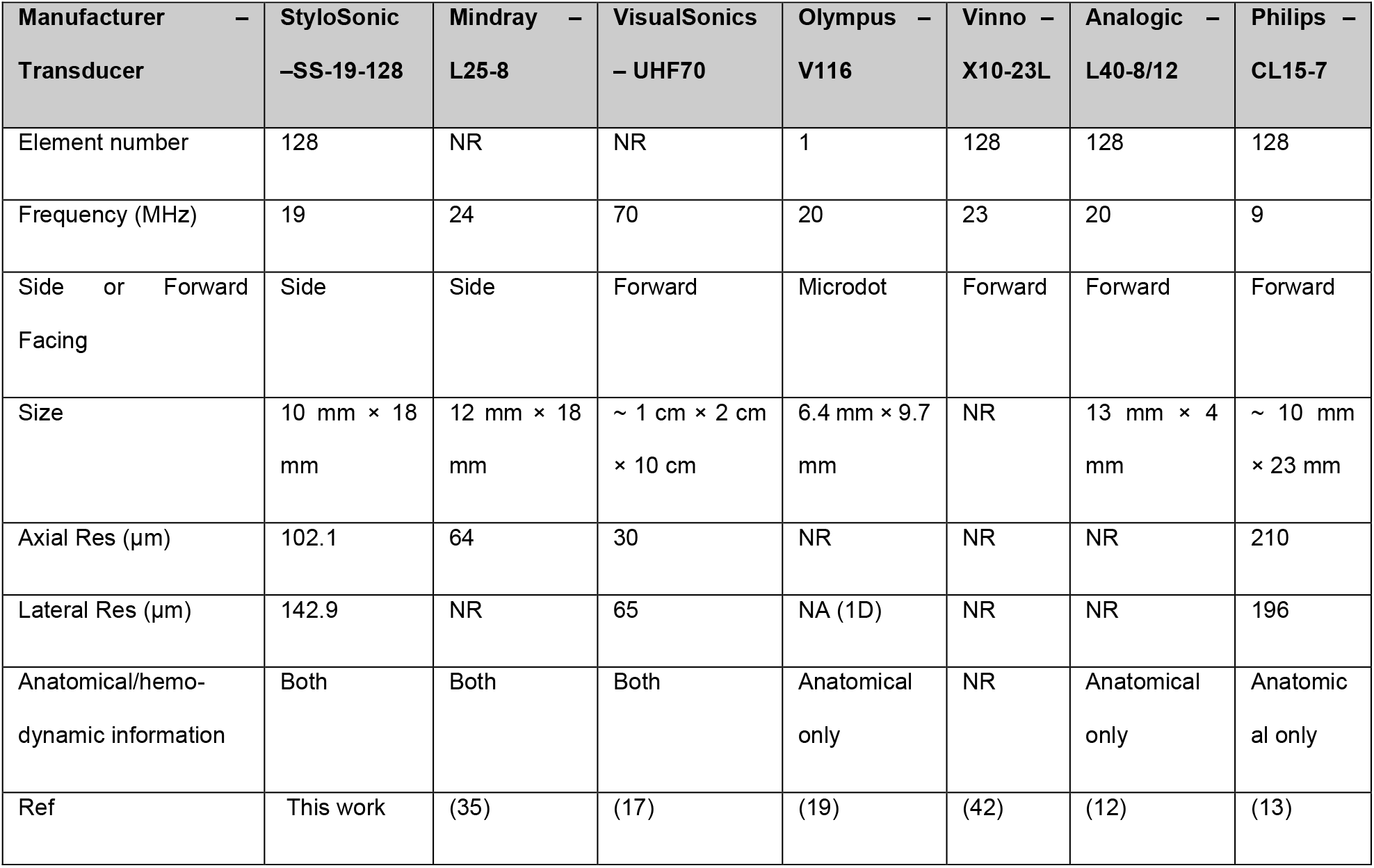
Comparison of similar transducers.

Element number, frequency, facing orientation, size, resolution and imaging mode of existing periodontal ultrasound transducers and our transducer are detailed in Table 1. The 128-element array and 19 MHz frequency of our transducer make it possible to image oral anatomies in a high resolution. The side facing design and miniaturized size guarantees its access to full mouth and the capacity to image all human teeth. The combination of B-mode imaging and Doppler imaging provides both anatomical and hemodynamic information in clinical applications.

The penetration of acoustic pressure waves decreases with increasing frequency due to faster attenuation.(43) The penetration depth of SS-19-128 was >30 mm, showing that besides periodontal structures, SS-19-128 can potentially achieve ultrasound tongue imaging (average tongue thickness is 41.3 mm in males and 39.3 mm in females).(44) Coherent compounding beamforming presented the highest SNR at all four depths from 15 mm to 30 mm. Coherent compounding had the best combination of resolution and SNR and was used for the subsequent work.

In the *in vivo* B-mode imaging, thanks to the high sensitivity to both hard and soft tissues of ultrasound, anatomical structures including gingival margin, AB, ABC, and CEJ were clearly visualized on all 28 teeth. By localizing these biomedical landmarks in ultrasonography, diagnosing and monitoring oral heath can be achieved in a fast, low-cost and non-invasive manner. Compared with the most recent work (13) that accessed 18 human teeth, SS-19-128 has 55.6% better range of access. The system was flexible and showed utility across a small cohort of five subjects.

Gingival thickness measurement results of SS-19-128 showed a good agreement with the gold standard - transgingival probing, which indicated the accuracy of the localizing the biomedical landmarks by SS-19-128 and its potential application in clinical oral health care.

The Doppler results supported that besides accessing anatomical structures by B-mode imaging, SS-19-128 can also provide comprehensive hemodynamic information to monitor oral health, which is significant in diagnosing periodontal diseases and evaluating surgery outcome, including tracking blood flow following soft tissue augmentation,(35) evaluating the degree of vascularization of oral tissues,(17) investigating pulpal microcirculation and vitality.(45)

In the future, we will expand the measurement to a larger population including healthy subjects and periodontitis patients. A power calculation was done to estimate how many teeth are needed with 80% power and an alpha of 0.05. We anticipate the bias to be ⩽ 0.20 mm based on our preliminary data. The power calculation suggests that a sample size of n=52 offers 80% power versus gold standard with a standard deviation of ⩽ 0.5 mm.

## CONCLUSION

This study reports the miniaturized SS-19-128 transducer to increase the accessible teeth range by 55.6% and offer full oral cavity assessment to monitor periodontal disease. SS-19-128 can easily access and image all human teeth including pre-molars and molars. Gingival thickness calibrated by ultrasound imaging strongly correlated with clinical methods (i.e., periodontal probing). Besides providing real-time imaging of the anatomical landmarks (i.e., CEJ, GM, ABC, etc.), real-time Doppler imaging to detect oral blood supply and hemodynamic information is also demonstrated on SS-19-128. Future work also includes improvement in excitation voltage and handpiece design. The highest voltage that can be applied to SS-19-128 is 5.6 V, which has an order of magnitude less than conventional transducers (30 V-50 V). Additional engineering of the handpiece will work to future reduce the form factor and increase ergonomics. Clinically, we will further expand the measurement to a large population including diseased subjects for clinical translation.

## Acknowledgements

A.H. and R.K.N. acknowledge federal funding from NIH under R43 DE031196. J.V.J. acknowledges NIH funding under R01 DE031307, R21 DE029025 and UL1 TR001442.

Some figures were prepared under a license with BioRender.

## References

1. Brennan DS, Spencer AJ, Roberts-Thomson KF. Tooth loss, chewing ability and quality of life. Qual Life Res. 2008;17(2):227–35.

2. Sischo L, Broder HL. Oral health-related quality of life: what, why, how, and future implications. J Dent Res. 2011;90(11):1264–70.

3. Eke PI, Dye BA, Wei L, Thornton-Evans GO, Genco RJ. Prevalence of periodontitis in adults in the United States: 2009 and 2010. J Dent Res. 2012;91(10):914–20.

4. Ghouth N, Duggal MS, BaniHani A, Nazzal H. The diagnostic accuracy of laser Doppler flowmetry in assessing pulp blood flow in permanent teeth: A systematic review. Dental Traumatology. 2018;34(5):311–9.

5. Grossi SG, Dunford RG, Ho A, Koch G, Machtei EE, Genco RJ. Sources of error for periodontal probing measurements. Journal of periodontal research. 1996;31(5):330–6.

6. Kumar V, Faizuddin M. Effect of smoking on gingival microvasculature: A histological study. J Indian Soc Periodontol. 2011;15(4):344–8.

7. Zoellner H, Chapple CC, Hunter N. Microvasculature in gingivitis and chronic periodontitis: disruption of vascular networks with protracted inflammation. Microsc Res Tech. 2002;56(1):15–31.

8. Orekhova LY, Barmasheva AA. Doppler flowmetry as a tool of predictive, preventive and personalised dentistry. EPMA J. 2013;4(1):21-.

9. Kripfgans OD, Chan H-L. Volumetric Ultrasound and Related Dental Applications. In: Chan H-L, Kripfgans OD, editors. Dental Ultrasound in Periodontology and Implantology: Examination, Diagnosis and Treatment Outcome Evaluation. Cham: Springer International Publishing; 2021. p. 231–43.

10. Chan H-LA, Kripfgans OD. Dental Ultrasound in Periodontology and Implantology.

11. Chifor R, Badea AF, Chifor I, Mitrea D-A, Crisan M, Badea ME. Periodontal evaluation using a non-invasive imaging method (ultrasonography). Medicine and Pharmacy Reports. 2019;92(Suppl No 3):S20–S32.

12. Nguyen K-CT, Duong DQ, Almeida FT, Major PW, Kaipatur NR, Pham T-T, et al. Alveolar bone segmentation in intraoral ultrasonographs with machine learning. J Dent Res. 2020;99(9):1054–61.

13. Fu L, Ling C, Jin Z, Luo J, Palma-Chavez J, Wu Z, et al. Photoacoustic imaging of posterior periodontal pocket using a commercial hockey-stick transducer. Journal of Biomedical Optics. 2022;27(5):056005.

14. Moore CA, Law JK, Retout M, Pham CT, Chang KCJ, Chen C, et al. High-resolution ultrasonography of gingival biomarkers for periodontal diagnosis in healthy and diseased subjects. Dentomaxillofac Radiol. 2022:20220044.

15. Mozaffarzadeh M, Moore C, Golmoghani EB, Mantri Y, Hariri A, Jorns A, et al. Motion-compensated noninvasive periodontal health monitoring using handheld and motor-based photoacoustic-ultrasound imaging systems. Biomedical Optics Express. 2021;12(3):1543–58.

16. Lin C, Chen F, Hariri A, Chen C, Wilder-Smith P, Takesh T, et al. Photoacoustic imaging for noninvasive periodontal probing depth measurements. J Dent Res. 2018;97(1):23–30.

17. Izzetti R, Vitali S, Aringhieri G, Oranges T, Dini V, Nisi M, et al. Discovering a new anatomy: exploration of oral mucosa with ultra-high frequency ultrasound. Dentomaxillofacial Radiology. 2020;49(7):20190318.

18. Sinjab K, Kripfgans OD, Ou A, Chan H-L. Ultrasonographic evaluation of edentulous crestal bone topography: A proof-of-principle retrospective study. Oral Surgery, Oral Medicine, Oral Pathology and Oral Radiology. 2022;133(1):110–7.

19. Nguyen K-CT, Le LH, Kaipatur NR, Major PW. Imaging the cemento-enamel junction using a 20-MHz ultrasonic transducer. Ultrasound in medicine & biology. 2016;42(1):333–8.

20. Chan HL, Wang HL, Fowlkes JB, Giannobile WV, Kripfgans OD. Non-ionizing real-time ultrasonography in implant and oral surgery: a feasibility study. Clinical oral implants research. 2017;28(3):341–7.

21. Nguyen K-CT, Le BM, Li M, Almeida FT, Major PW, Kaipatur NR, et al. Localization of cementoenamel junction in intraoral ultrasonographs with machine learning. Journal of Dentistry. 2021;112:103752.

22. Pan Y-C, Chan H-L, Kong X, Hadjiiski LM, Kripfgans OD. Multi-class deep learning segmentation and automated measurements in periodontal sonograms of a porcine model. Dentomaxillofacial Radiology. 2021;50:20210363.

23. Yi J, Nguyen K-CT, Wang W, Yang W, Pan M, Lou E, et al. Polyacrylamide/Alginate double-network tough hydrogels for intraoral ultrasound imaging. Journal of colloid and interface science. 2020;578:598–607.

24. Chifor R, Li M, Nguyen K-CT, Arsenescu T, Chifor I, Badea AF, et al. Three-dimensional periodontal investigations using a prototype handheld ultrasound scanner with spatial positioning reading sensor. Medical Ultrasonography. 2021;23(3):297–304.

25. Palma-Chavez J, Wear KA, Mantri Y, Jokerst JV, Vogt WC. Photoacoustic imaging phantoms for assessment of object detectability and boundary buildup artifacts. Photoacoustics. 2022;26:100348.

26. Matrone G, Savoia AS, Caliano G, Magenes G, editors. Ultrasound plane-wave imaging with delay multiply and sum beamforming and coherent compounding. 2016 38th Annual International Conference of the IEEE Engineering in Medicine and Biology Society (EMBC); 2016: IEEE.

27. Huang L, Huang Y, Gao K, editors. Transrectal ultrasound imaging using plane-wave, fan-beam and wide-beam ultrasound: Phantom results. Medical Imaging 2019: Physics of Medical Imaging; 2019: SPIE.

28. Le DQ, Chang E, Dayton PA, Johnson K, editors. High-framerate dynamic contrast-enhanced ultrasound imaging of rat kidney perfusion. 2019 IEEE International Ultrasonics Symposium (IUS); 2019: IEEE.

29. Siesky BA, Harris A, Kagemann L. Chapter 10 - Imaging of Ocular Blood Flow. In: Huang D, Kaiser PK, Lowder CY, Traboulsi EI, editors. Retinal Imaging. Philadelphia: Mosby; 2006. p. 134–41.

30. Slak B, Daabous A, Bednarz W, Strumban E, Maev RG. Assessment of gingival thickness using an ultrasonic dental system prototype: A comparison to traditional methods. Annals of Anatomy-Anatomischer Anzeiger. 2015;199:98–103.

31. Vandana K, Savitha B. Thickness of gingiva in association with age, gender and dental arch location. Journal of clinical periodontology. 2005;32(7):828–30.

32. Kloukos D, Koukos G, Doulis I, Sculean A, Stavropoulos A, Katsaros C. Gingival thickness assessment at the mandibular incisors with four methods: A cross-sectional study. Journal of periodontology. 2018;89(11):1300–9.

33. Schneider CA, Rasband WS, Eliceiri KW. NIH Image to ImageJ: 25 years of image analysis. Nature methods. 2012;9(7):671–5.

34. Rossmann K. Point spread-function, line spread-function, and modulation transfer function: tools for the study of imaging systems. Radiology. 1969;93(2):257–72.

35. Tavelli L, Barootchi S, Majzoub J, Chan HL, Giannobile WV, Wang HL, et al. Ultrasonographic tissue perfusion analysis at implant and palatal donor sites following soft tissue augmentation: A clinical pilot study. Journal of Clinical Periodontology. 2021;48(4):602–14.

36. Perry DA, Beemsterboer P, Essex G. Periodontology for the dental hygienist: Saunders Elsevier; 2007.

37. Chan H-LA, Kripfgans OD. Ultrasonic Imaging for Evaluating Peri-Implant Diseases. Dental Ultrasound in Periodontology and Implantology: Springer; 2021. p. 161–75.

38. Aldrich JE. Basic physics of ultrasound imaging. Critical care medicine. 2007;35(5):S131–S7.

39. Ng A, Swanevelder J. Resolution in ultrasound imaging. Continuing Education in Anaesthesia Critical Care & Pain. 2011;11(5):186–92.

40. Rothlübbers S, Strohm H, Eickel K, Jenne J, Kuhlen V, Sinden D, et al., editors. Improving image quality of single plane wave ultrasound via deep learning based channel compounding. 2020 IEEE International Ultrasonics Symposium (IUS); 2020: IEEE.

41. Chen Z, Zhong J, Ouyang X, Zhou S, Xie Y, Lou X. Gingival thickness assessment of gingival recession teeth. Beijing da xue xue bao Yi xue ban= Journal of Peking University Health Sciences. 2020;52(2):339–45.

42. https://www.vinno-ultraschall.de/vinno-x10-23l/ [

43. Bader KB, Crowe MJ, Raymond JL, Holland CK. Effect of Frequency-Dependent Attenuation on Predicted Histotripsy Waveforms in Tissue-Mimicking Phantoms. Ultrasound in medicine & biology. 2016;42(7):1701–5.

44. Nakamori M, Imamura E, Fukuta M, Tachiyama K, Kamimura T, Hayashi Y, et al. Tongue thickness measured by ultrasonography is associated with tongue pressure in the Japanese elderly. PloS one. 2020;15(8):e0230224.

45. Chen E, Abbott PV. Dental pulp testing: a review. International journal of dentistry. 2009;2009.

